# Preliminary results of models to predict areas in the Americas with increased likelihood of Zika virus transmission in 2017

**DOI:** 10.1101/187591

**Authors:** The ZIKAVAT Collaboration, Jason Asher, Christopher Barker, Grace Chen, Derek Cummings, Matteo Chinazzi, Shelby Daniel-Wayman, Marc Fischer, Neil Ferguson, Dean Follman, M. Elizabeth Halloran, Michael Johansson, Kiersten Kugeler, Jennifer Kwan, Justin Lessler, Ira M. Longini, Stefano Merler, Andrew Monaghan, Ana Pastore y Piontti, Alex Perkins, D. Rebecca Prevots, Robert Reiner, Luca Rossi, Isabel Rodriguez-Barraquer, Amir S. Siraj, Kaiyuan Sun, Alessandro Vespignani, Qian Zhang

**Affiliations:** Biomedical Advanced Research and Development Authority, Washington, DC, USA; University of California, Davis, California, USA; National Institutes of Health, Bethesda, MD,USA; Johns Hopkins, Bloomberg School of Public Health, Baltimore, Maryland, USA; University of Florida, Gainesville, Florida, USA; Northeastern University, Boston, Massachusetts, USA; Centers for Disease Control and Prevention, Fort Collins, Colorado and San Juan, Puerto Rico, USA; Imperial College, London, UK; Fred Hutchinson Center, Seattle,WA, USA; University of Washington, Seattle, Washington, USA; Bruno Kessler Foundation, Trento, Italy; National Center for Atmospheric Research, Boulder, Colorado, USA; University of Notre Dame, Notre Dame, Indiana, USA; Institute for Scientific Interchange Foundation, Turin, Italy; University of California, San Francisco, California, USA

**Author notes:** Corresponding author: Marc Fischer, Arboviral Diseases Branch, Centers for Disease Control and Prevention, Fort Collins, Colorado,. Zika Modeling and Projections for Vaccination Trials Collaboration.

## Abstract

Numerous Zika virus vaccines are being developed. However, identifying sites to evaluate the efficacy of a Zika virus vaccine is challenging due to the general decrease in Zika virus activity. We compare results from three different modeling approaches to estimate areas that may have increased relative risk of Zika virus transmission during 2017. The analysis focused on eight priority countries (i.e., Brazil, Colombia, Costa Rica, Dominican Republic, Ecuador, Mexico, Panama, and Peru). The models projected low incidence rates during 2017 for all locations in the priority countries but identified several subnational areas that may have increased relative risk of Zika virus transmission in 2017. Given the projected low incidence of disease, the total number of participants, number of study sites, or duration of study follow-up may need to be increased to meet the efficacy study endpoints.

## Introduction

Zika virus is a mosquito-borne flavivirus primarily transmitted to humans by *Aedes (Stegomyia)* species mosquitoes [Petersen 2016]. The virus was first identified in Uganda in 1947 [Dick 1952]. Prior to 2007, only sporadic human disease cases were reported from countries in Africa and Asia. From 2007-2014, outbreaks were identified in Southeast Asia and the Western Pacific [Duffy 2009; Heang 2012; Cao-Lormeau 2013; Roth 2014]. In 2015, Zika virus was identified for the first time in the Americas with large outbreaks reported in Brazil and subsequent spread throughout the region [Zanluca 2015; Ikejezie 2017].

Most Zika virus infections are asymptomatic [Duffy 2009]. For patients with symptomatic illness, disease is generally mild and characterized by acute onset of fever or rash. However, Zika virus infection during pregnancy can cause adverse outcomes such as fetal loss, congenital microcephaly, and other serious birth defects [Moore 2017; Rasmussen 2016]. There are no vaccines to prevent Zika virus infection. However, numerous candidate vaccines are being developed and several have entered clinical trials [Thomas 2017].

The National Institutes of Health (NIH) Vaccine Research Center is conducting a Phase 2B clinical trial to evaluate the safety, immunogenicity, and efficacy of a Zika virus DNA vaccine in healthy adolescents and adults. The study will begin in July 2017 and will be performed at multiple sites in the Americas. The current protocol proposes to enroll 2,400 subjects randomized on a 1:1 basis to receive the study vaccine or placebo. Assuming a 50% vaccine efficacy, the study could be completed in approximately 2 years if the average annual incidence of symptomatic Zika virus disease among participants receiving placebo is ≥2%. The sample size or study duration will need to be increased if the symptomatic disease rate among participants is <2% or if >10% of the participants are already protected at baseline.

Many factors impact the likelihood and rate of ongoing Zika virus infections in a population (e.g., presence and abundance of vector mosquitoes, temperature, precipitation, human mobility, population density, living conditions, and baseline immunity). However, there are limited data and experience for predicting the occurrence and magnitude of future Zika virus disease outbreaks in the Americas. To help with study site selection to meet the efficacy endpoint, NIH and the Centers for Disease Control and Prevention (CDC) requested assistance from three academic groups to adapt and apply existing mathematical models to estimate areas that may have increased likelihood of Zika virus transmission in 2017. Comparing results from three different modeling approaches enables better characterization of the predictive uncertainty due to model and data limitations with higher confidence assigned to predictions for areas where model agreement is strong. This report synthesizes findings from the three models obtained in early 2017.

## Methods

We identified three models that had been developed and used to predict the geographic location or incidence of dengue or Zika virus disease in 2015-2016. Each modeling team was tasked with providing a list of areas in the Americas with the highest probability of Zika virus transmission and estimated infection rates during 2017. Each model used different input variables and output measures, and relied on different units of reporting. Therefore, comparisons were limited to subnational areas within the same countries rather than between countries.

To facilitate comparisons between models, data were aggregated to a state/province level and focused on eight priority countries (i.e., Brazil, Colombia, Costa Rica, Dominican Republic, Ecuador, Mexico, Panama, and Peru). These countries were selected based on their capacity and infrastructure to perform clinical trials, and preliminary assessments of expected Zika virus activity based on surveillance reports and previous experience with dengue and chikungunya viruses.

### Modeling Team 1 (MT1)^1^

The Global Epidemic and Mobility Model (GLEAM) is a discrete stochastic epidemic computational model based on a meta-population approach in which the world is defined in geographical census areas connected in a network of interactions by human travel fluxes corresponding to transportation infrastructures and mobility patterns [Zhang 2017]. The model includes a multiscale mobility model integrating different layers of transportation networks ranging from long-range airline connections to short-range daily commuting patterns. GLEAM also integrates high-resolution demographic, socioeconomic, temperature, and vector abundance data. The model has been used to analyze the spatiotemporal spread and magnitude of the Zika epidemic in the Americas accounting for seasonal environmental factors and detailed population data. The model is fully stochastic and from any nominally identical initialization (initial conditions and disease model) generates an ensemble of possible epidemic evolutions for epidemic observables, such as newly generated cases, time of arrival of the infection, and number of traveling carriers. The model native grid cell resolution is 25 km x 25 km and cells are aggregated/projected to the desired level of resolution. For the purpose of studying the Zika outbreak, the model outputs include: 1) the median projected infection rate and 95% confidence intervals (95% CI) for each state/province in the eight priority countries (**Tables 1–8**); and 2) the probability that an urban area in any country/territory in the Americas will experience an annual Zika virus infection rate ≥10% in 2017 (**Table 9**).

**Table 1:** Brazil modeling results by state.

| State | Median Rank | MT1 Rank | MT2 Rank | MT3 Rank | Population | MT1* Infection rate | (95% CI) | MT2† Susc. pop | (95%CI) | MT3 Score |
| --- | --- | --- | --- | --- | --- | --- | --- | --- | --- | --- |
| Minas Gerais | 1 | 1 | 19 | 1 | 19,987,031 | 0.06% | (0.00-0.35) | 13.2% | (6.8-21.5) | 0.023 |
| São Paulo | 3 | 2 | 26 | 3 | 41,315,532 | 0.01% | (0.00-0.03) | 6.0% | (4.1-13.3) | 0.106 |
| Maranhão | 5 | 12 | 5 | 5 | 6,401,099 | <0.01% |  | 32.1% | (29.1-35.3) | 0.224 |
| Amazonas | 8 | 12 | 3 | -- | 3,585,205 | <0.01% |  | 33.9% | (27.2-40.2) | -- |
| Ceará | 8 | 12 | 4 | 8 | 8,382,131 | <0.01% |  | 33.5% | (30.2-36.4) | 0.316 |
| Mato Grosso | 9 | 3 | 27 | 9 | 3,060,605 | <0.01% | (0.00-0.07) | 3.2% | (-29.6-20.1) | 0.443 |
| Mato Grosso do Sul | 9 | 9 | 7 | 16 | 2,506,342 | <0.01% | (0.00-0.52) | 30.2% | (27.0-34.5) | 0.676 |
| Pará | 10 | 12 | 10 | 2 | 7,438,518 | <0.01% |  | 27.3% | (25.1-30.3) | 0.03 |
| Rio Grande do Sul | 10 | 10 | 22 | 6 | 10,482,802 | <0.01% | (0.00-0.03) | 11.5% | (4.5-19.9) | 0.268 |
| Rondônia | 11 | 11 | 17 | 4 | 1,576,505 | <0.01% | (0.00-0.02) | 22.5% | (18.5-26.3) | 0.124 |
| Rio de Janeiro | 11 | 5 | 21 | 11 | 14,324,781 | <0.01% | (0.00-0.03) | 11.8% | (-4.2-20.6) | 0.502 |
| Rio Grande do Norte | 12 | 12 | 6 | 20 | 3,023,570 | <0.01% |  | 31.5% | (26.5-35.2) | 0.799 |
| Roraima | 12 | 12 | 1 | 23 | 456,864 | <0.01% |  | 37.9% | (31.3-44.1) | 0.962 |
| Piauí | 12 | 12 | 2 | 15 | 3,377,661 | <0.01% |  | 36.2% | (33.9-39.3) | 0.666 |
| Tocantins | 12 | 12 | 14 | 7 | 1,435,936 | <0.01% |  | 24.6% | (14.7-30.5) | 0.275 |
| Distrito Federal | 12 | 12 | 23 | 10 | 2,647,348 | <0.01% |  | 7.4% | (0.9-18.1) | 0.448 |
| Acre | 12 | 12 | 13 | 12 | 708,021 | <0.01% |  | 26.2% | (20.1-30.2) | 0.522 |
| Pernambuco | 12 | 12 | 9 | 19 | 7,766,940 | <0.01% |  | 29.1% | (26.2-32.5) | 0.79 |
| Alagoas | 12 | 8 | 12 | 18 | 2,812,590 | <0.01% | (0.00-0.01) | 26.7% | (19.3-31.6) | 0.761 |
| Sergipe | 12 | 12 | 8 | 21 | 1,991,960 | <0.01% |  | 29.9% | (27.3-32.3) | 0.883 |
| Paraíba | 12 | 12 | 11 | 22 | 3,619,622 | <0.01% |  | 27.1% | (22.5-31.4) | 0.948 |
| Goiás | 13 | 12 | 16 | 13 | 6,205,103 | <0.01% |  | 23.9% | (18.8-28.8) | 0.53 |
| Bahia | 14 | 12 | 20 | 14 | 12,687,903 | <0.01% |  | 12.1% | (-6.2-22.3) | 0.66 |
| Espírito Santo | 15 | 4 | 15 | 17 | 2,682,726 | <0.01% | (0.00-0.01) | 24.4% | (19.9-28.3) | 0.735 |
| Amapá | 15 | 12 | 18 | -- | 594,387 | <0.01% |  | 20.7% | (17.7-22.8) | -- |
| Santa Catarina | 16 | 7 | 24 | -- | 5,417,796 | <0.01% | (0.00-0.34) | 7.3% | (2.8-16.1) | -- |
| Paraná | 16 | 6 | 25 | -- | 10,616,043 | <0.01% | (0.00-0.63) | 6.8% | (4.3-14.8) | -- |

\*Median and 95% confidence intervals for projected infection rates in 2017 from >1,000 simulations.
† Proportion of the population still at risk for infection.

**Table 2:** Colombia modeling results by state.

| State | Median Rank | MT1 Rank | MT2 Rank | MT3 Rank | Population | MT1* Inf. rate | (95% CI) | MT2† Susc. pop | (95%CI) | MT3 Score |
| --- | --- | --- | --- | --- | --- | --- | --- | --- | --- | --- |
| Nariño | 2 | 2 | 19 | 2 | 1,766,008 | 0.50% | (0.04-2.17) | 2.7% | (1.6-5.7) | 0.074 |
| La Guajira | 3 | 1 | 3 | 4 | 985,498 | 0.58% | (0.01-3.46) | 30.2% | (24.1-35.9) | 0.137 |
| Córdoba | 4 | 4 | 1 | 11 | 1,736,218 | 0.26% | (0.02-1.00) | 33.5% | (30.0-37.1) | 0.301 |
| Bolívar | 5 | 9 | 4 | 5 | 2,122,021 | 0.05% | (0.00-0.23) | 29.8% | (23.8-34.4) | 0.157 |
| Sucre | 7 | 5 | 7 | 7 | 859,909 | 0.16% | (0.01-0.77) | 27.3% | (17.3-33.5) | 0.205 |
| Antioquia | 9 | 8 | 18 | 9 | 6,534,764 | 0.06% | (0.01-0.39) | 3.7% | (1.0-10.6) | 0.227 |
| Guainía | 9 | 13 | 5 | -- | 42,123 | 0.01% | (0.00-16.07) | 28.8% | (25.3-32.0) | -- |
| Cauca | 10 | 10 | 15 | 3 | 1,391,889 | 0.05% | (0.00-0.30) | 7.8% | (5.6-10.2) | 0.135 |
| Magdalena | 11 | 6 | 11 | 16 | 1,272,278 | 0.16% | (0.02-0.70) | 18.9% | (5.0-26.6) | 0.561 |
| Cesar | 11 | 11 | 9 | 12 | 1,041,203 | 0.02% | (0.00-0.18) | 26.7% | (17.6-32.8) | 0.330 |
| Vichada | 12 | 12 | 2 | 17 | 73,702 | 0.02% | (0.00-10.75) | 31.9% | (24.3-37.9) | 0.620 |
| Choco | 12 | 23 | 12 | 8 | 505,046 | <0.01% |  | 13.5% | (5.4-21.9) | 0.222 |
| Caquetá | 13 | 7 | 13 | 19 | 483,834 | 0.06% | (0.01-0.36) | 9.5% | (-4.0-17.5) | 0.641 |
| Risaralda | 14 | 14 | 22 | 13 | 957,250 | <0.01% | (0.00-0.03) | 0.1% | (-8.4-9.6) | 0.370 |
| Vaupés | 14 | 17 | 10 | -- | 44,079 | <0.01% | (0.00-1.43) | 23.8% | (17.4-27.6) | -- |
| Atlántico | 14 | 20 | 8 | 14 | 2,489,709 | <0.01% | (0.00-0.01) | 27.1% | (12.1-37.0) | 0.404 |
| Caldas | 15 | 15 | 20 | 1 | 989,942 | <0.01% | (0.00-0.02) | 1.6% | (-0.8-5.7) | 0.068 |
| Putumayo | 16 | 3 | 16 | 18 | 349,537 | 0.38% | (0.03-3.02) | 7.3% | (-5.1-16.5) | 0.632 |
| Guaviare | 16 | 16 | 6 | 21 | 112,621 | <0.01% | (0.00-0.19) | 28.6% | (17.2-36.4) | 0.774 |
| Boyacá | 19 | 19 | 21 | 6 | 1,278,061 | <0.01% | (0.00-0.01) | 1.2% | (-0.3-2.1) | 0.169 |
| Amazonas | 19 | 23 | 14 | -- | 77,088 | <0.01% |  | 9.0% | (-18.3-22.2) | -- |
| Meta | 20 | 21 | 17 | 20 | 979,683 | <0.01% | (0.00-0.01) | 5.4% | (-19.1-17.2) | 0.768 |
| Cundinamarca | 23 | 23 | 24 | 15 | 2,721,368 | <0.01% |  | -0.6% | (-3.3-0.6) | 0.496 |
| Quindío | 23 | 23 | 23 | 10 | 568,473 | <0.01% |  | -0.1% | (-4.6-11.2) | 0.235 |
| Santander | 23 | 23 | 27 | 23 | 2,071,044 | <0.01% |  | -6.6% | (-32.9-6.3) | 0.829 |
| Tolima | 23 | 23 | 25 | 22 | 1,412,230 | <0.01% |  | -1.5% | (-29.1-11.5) | 0.793 |
| Huila | 24 | 18 | 29 | 24 | 1,168,910 | <0.01% | (0.00-0.01) | -9.5% | (-41.2-5.2) | 0.832 |
| Norte Santander | 25 | 22 | 31 | 25 | 1,367,716 | <0.01% | (0.00-0.01) | -19.8% | (-71.1-4.3) | 0.837 |
| Valle | 26 | 23 | 26 | 26 | 4,660,438 | <0.01% |  | -5.2% | (-38.8-11.1) | 0.871 |
| Arauca | 27 | 23 | 28 | 27 | 265,190 | <0.01% |  | -7.1% | (-44.2-11.3) | 0.962 |
| Casanare | 28 | 23 | 30 | 28 | 362,698 | <0.01% |  | -16.7% | (-73.6-11.9) | 0.993 |

\*Median and 95% confidence intervals for projected infection rates in 2017 from >1,000 simulations.
† Proportion of the population still at risk for infection.

**Table 3:** Costa Rica modeling results by province.

| Province | Median Rank | MT1 Rank | MT2 Rank | MT3 Rank | Population | MT1*<br>Inf. rate | (95% CI) | MT2†<br>Susc.<br>pop | (95%CI) | MT3<br>Score |
| --- | --- | --- | --- | --- | --- | --- | --- | --- | --- | --- |
| Limon | 3 | 2 | 4 | -- | 386,862 | <0.01% | (0.00-3.99) | 3.0% | (-0.7-13.8) |  |
| Puntarenas | 3 | 3 | 2 | -- | 410,929 | <0.01% | (0.00-1.03) | 7.8% | (-0.6-15.7) |  |
| Guanacaste | 3 | 5 | 1 | -- | 354,154 | <0.01% |  | 16.5% | (10.6-22.1) |  |
| Cartago | 4 | 1 | 7 | -- | 490,903 | <0.01% | (0.00-1.51) | 0.8% | (0.2-3.5) |  |
| Alajuela | 4 | 5 | 3 | -- | 885,571 | <0.01% |  | 6.1% | (3.1-15.3) |  |
| San Jose | 5 | 4 | 6 | -- | 1,404,242 | <0.01% | (0.00-0.38) | 1.3% | (0.1-7.8) |  |
| Heredia | 5 | 5 | 5 | -- | 433,677 | <0.01% |  | 1.8% | (0.5-10.1) |  |

\*Median and 95% confidence intervals for projected infection rates in 2017 from >1,000 simulations.
† Proportion of the population still at risk for infection.

**Table 4:**
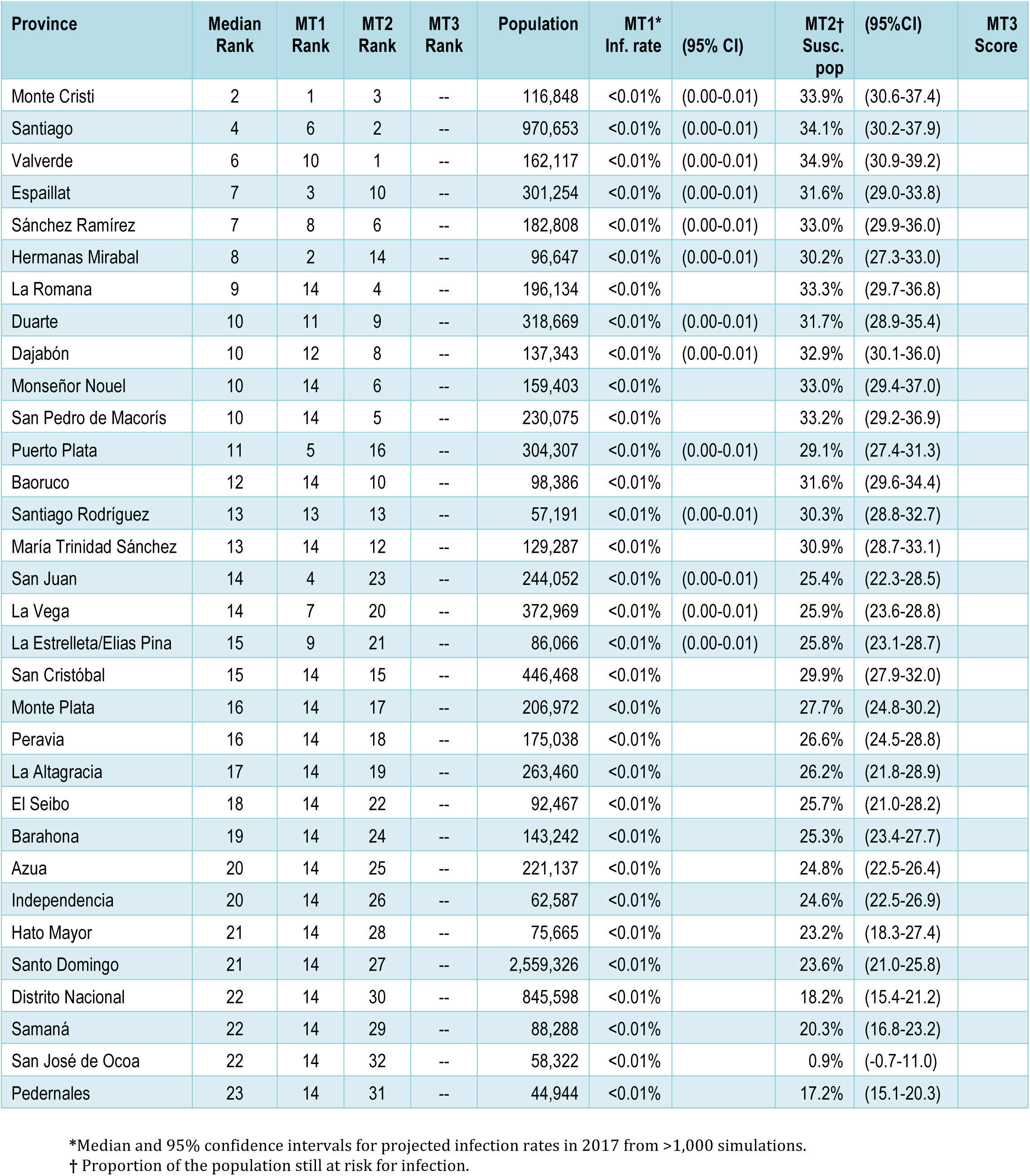
Dominican Republic modeling results by province.

**Table 5:** Ecuador modeling results by province.

| Province | Median Rank | MT1 Rank | MT2 Rank | MT3 Rank | Population | MT1* Inf. rate | (95% CI) | MT2† Susc. pop | (95%CI) | MT3 Score |
| --- | --- | --- | --- | --- | --- | --- | --- | --- | --- | --- |
| Sucumbíos | 4 | 1 | 6 | -- | 186,504 | 7.55% | (0.21-15.90) | 14.8% | (6.6-24.4) |  |
| Los Ríos | 5 | 5 | 4 | -- | 777,079 | 0.81% | (0.07-3.59) | 18.6% | (12.9-24.9) |  |
| Esmeraldas | 5 | 2 | 8 | -- | 555,848 | 5.81% | (0.43-12.96) | 12.0% | (5.3-21.2) |  |
| Orellana | 6 | 3 | 9 | -- | 88,611 | 5.43% | (0.38-12.54) | 8.6% | (4.1-19.9) |  |
| Manabí | 7 | 6 | 7 | -- | 1,281,663 | 0.55% | (0.05-1.83) | 13.0% | (2.5-21.3) |  |
| Guayas | 7 | 11 | 2 | -- | 3,595,034 | 0.02% | (0.00-0.13) | 27.7% | (24.4-30.4) |  |
| El Oro | 8 | 13 | 3 | -- | 613,666 | <0.01% | (0.00-0.62) | 23.0% | (18.9-27.3) |  |
| Santa Elena | 8 | 15 | 1 | -- | 292,220 | <0.01% | (0.00-0.01) | 30.0% | (27.0-32.8) |  |
| Pastaza | 9 | 4 | 14 | -- | 87,380 | 3.07% | (0.49-6.55) | 2.8% | (1.1-12.3) |  |
| Bolívar | 12 | 7 | 16 | -- | 200,410 | 0.51% | (0.04-2.36) | 1.4% | (0.3-6.3) |  |
| Napo | 12 | 8 | 16 | -- | 106,649 | 0.48% | (0.06-1.30) | 1.4% | (0.2-10.8) |  |
| Morona Santiago | 12 | 12 | 12 | -- | 153,622 | 0.02% | (0.00-0.31) | 5.6% | (2.2-16.2) |  |
| Galápagos | 12 | 19 | 5 | -- | 20,541 | <0.01% |  | 16.6% | (10.6-21.6) |  |
| Cotopaxi | 14 | 10 | 18 | -- | 394,132 | 0.24% | (0.02-1.21) | 0.7% | (0.1-2.7) |  |
| Cañar | 14 | 17 | 10 | -- | 234,668 | <0.01% | (0.00-0.02) | 6.9% | (5.0-8.7) |  |
| Carchi | 15 | 9 | 20 | -- | 155,648 | 0.34% | (0.00-1.64) | 0.0% | (0.0-0.3) |  |
| Zamora Chinchipe | 15 | 14 | 15 | -- | 105,214 | <0.01% | (0.00-0.24) | 2.2% | (0.2-13.3) |  |
| Loja | 15 | 16 | 13 | -- | 450,634 | <0.01% | (0.00-0.17) | 3.6% | (2.3-7.6) |  |
| Santo Domingo Tsáchilas | 15 | 19 | 11 | -- | 422,080 | <0.01% |  | 6.0% | (1.6-17.8) |  |
| Azuay | 18 | 18 | 18 | -- | 726,744 | <0.01% | (0.00-0.03) | 0.7% | (0.5-1.1) |  |
| Chimborazo | 20 | 19 | 20 | -- | 468,077 | <0.01% |  | 0.0% | (0.0-0.2) |  |
| Imbabura | 20 | 19 | 20 | -- | 394,164 | <0.01% |  | 0.0% | (0.0-0.3) |  |
| Pichincha | 20 | 19 | 20 | -- | 2,449,094 | <0.01% |  | 0.0% | (-0.1-0.3) |  |
| Tungurahua | 20 | 19 | 20 | -- | 530,294 | <0.01% |  | 0.0% | (0.0-0.0) |  |

\*Median and 95% confidence intervals for projected infection rates in 2017 from >1,000 simulations.
† Proportion of the population still at risk for infection.

**Table 6:** Mexico modeling results by state

| State | Median Rank | MT1 Rank | MT2 Rank | MT3 Rank | Population | MT1* Inf. rate | (95% CI) | MT2† Susc. pop | (95%CI) | MT3 Score |
| --- | --- | --- | --- | --- | --- | --- | --- | --- | --- | --- |
| Sinaloa | 1 | 1 | 1 | 4 | 2,838,630 | 2.44% | (0.02-14.95) | 38.5% | (31.2-44.1) | 0.166 |
| San Luis Potosí | 4 | 4 | 19 | 1 | 2,626,925 | 0.33% | (0.00-4.29) | 10.0% | (9.0-10.9) | 0.002 |
| Tamaulipas | 5 | 5 | 2 | 7 | 2,684,165 | 0.28% | (0.00-3.86) | 37.4% | (29.7-42.6) | 0.286 |
| Sonora | 9 | 3 | 9 | 14 | 2,663,174 | 0.58% | (0.03-1.65) | 29.1% | (25.5-32.8) | 0.552 |
| Quintana Roo | 10 | 10 | 7 | 19 | 1,127,393 | 0.03% | (0.00-1.56) | 30.6% | (25.5-34.1) | 0.695 |
| Baja California Sur | 11 | 2 | 11 | 17 | 569,628 | 1.55% | (0.12-4.85) | 25.4% | (22.4-28.4) | 0.622 |
| Nayarit | 11 | 8 | 12 | 11 | 10,443,171 | 0.05% | (0.00-11.01) | 24.7% | (20.6-27.7) | 0.405 |
| Campeche | 12 | 12 | 6 | 22 | 799,030 | 0.02% | (0.00-1.29) | 30.9% | (24.9-35.2) | 0.797 |
| Nuevo León | 12 | 21 | 3 | 12 | 4,881,286 | <0.01% | (0.00-0.29) | 35.1% | (27.5-40.1) | 0.468 |
| Veracruz | 14 | 6 | 14 | 26 | 7,620,285 | 0.18% | (0.00-4.59) | 21.5% | (19.8-22.9) | 0.936 |
| Guerrero | 14 | 14 | 10 | 21 | 3,479,330 | 0.01% | (0.00-2.08) | 26.8% | (24.8-28.8) | 0.754 |
| Chihuahua | 15 | 13 | 17 | -- | 3,309,967 | 0.01% | (0.00-0.06) | 15.4% | (8.9-18.9) | -- |
| Colima | 15 | 15 | 7 | 25 | 671,505 | 0.01% | (0.00-18.45) | 30.6% | (26.4-34.1) | 0.846 |
| Coahuila | 15 | 25 | 15 | 15 | 3,197,854 | <0.01% | (0.00-0.01) | 20.0% | (15.4-22.9) | 0.614 |
| Chiapas | 16 | 9 | 16 | 16 | 4,960,867 | 0.04% | (0.00-2.50) | 15.8% | (14.0-20.1) | 0.615 |
| Jalisco | 16 | 16 | 21 | 8 | 7,708,204 | 0.01% | (0.00-1.61) | 6.4% | (3.2-18.2) | 0.331 |
| Michoacán | 17 | 17 | 20 | 2 | 4,448,339 | 0.01% | (0.00-6.27) | 6.8% | (6.2-9.2) | 0.023 |
| Oaxaca | 18 | 7 | 18 | 20 | 3,846,384 | 0.11% | (0.00-5.65) | 13.0% | (11.5-17.5) | 0.697 |
| Hidalgo | 18 | 11 | 24 | 18 | 3,058,115 | 0.02% | (0.00-0.42) | 3.6% | (2.7-4.5) | 0.651 |
| Durango | 18 | 18 | 22 | 6 | 1,315,740 | <0.01% | (0.00-1.24) | 4.8% | (3.4-15.0) | 0.280 |
| Tabasco | 19 | 19 | 5 | 23 | 2,301,101 | <0.01% | (0.00-0.80) | 32.3% | (28.9-35.7) | 0.808 |
| Puebla | 20 | 20 | 25 | 9 | 6,319,334 | <0.01% | (0.00-0.73) | 2.8% | (2.3-4.3) | 0.354 |
| Yucatán | 22 | 22 | 4 | 27 | 2,018,565 | <0.01% | (0.00-0.04) | 33.3% | (28.1-37.4) | 0.957 |
| Zacatecas | 23 | 23 | 28 | 13 | 1,582,975 | <0.01% | (0.00-0.06) | 0.6% | (0.3-3.6) | 0.507 |
| Guanajuato | 23 | 24 | 23 | 10 | 5,825,418 | <0.01% | (0.00-0.07) | 4.1% | (0.5-15.1) | 0.382 |
| Morelos | 24 | 26 | 13 | 24 | 1,876,080 | <0.01% |  | 23.2% | (19.2-27.4) | 0.837 |
| Baja California | 26 | 26 | 26 | -- | 3,059,921 | <0.01% |  | 1.5% | (0.8-10.2) | -- |
| México | 26 | 26 | 30 | 3 | 17,214,830 | <0.01% |  | 0.2% | (0.2-0.4) | 0.143 |
| Querétaro | 26 | 26 | 27 | 5 | 1,932,829 | <0.01% |  | 0.9% | (0.7-4.2) | 0.272 |
| Aguascalientes | 28 | 26 | 29 | -- | 1,206,312 | <0.01% |  | 0.5% | (0.0-12.9) |  |
| Distrito Federal | 29 | 26 | 31 | -- | 7,924,645 | <0.01% |  | 0.0% | (0.0-0.0) |  |
| Tlaxcala | 29 | 26 | 31 | -- | 1,066,673 | <0.01% |  | 0.0% | (0.0-0.0) |  |

\*Median and 95% confidence intervals for projected infection rates in 2017 from > 1,000 simulations.
† Proportion of the population still at risk for infection.

**Table 7:**
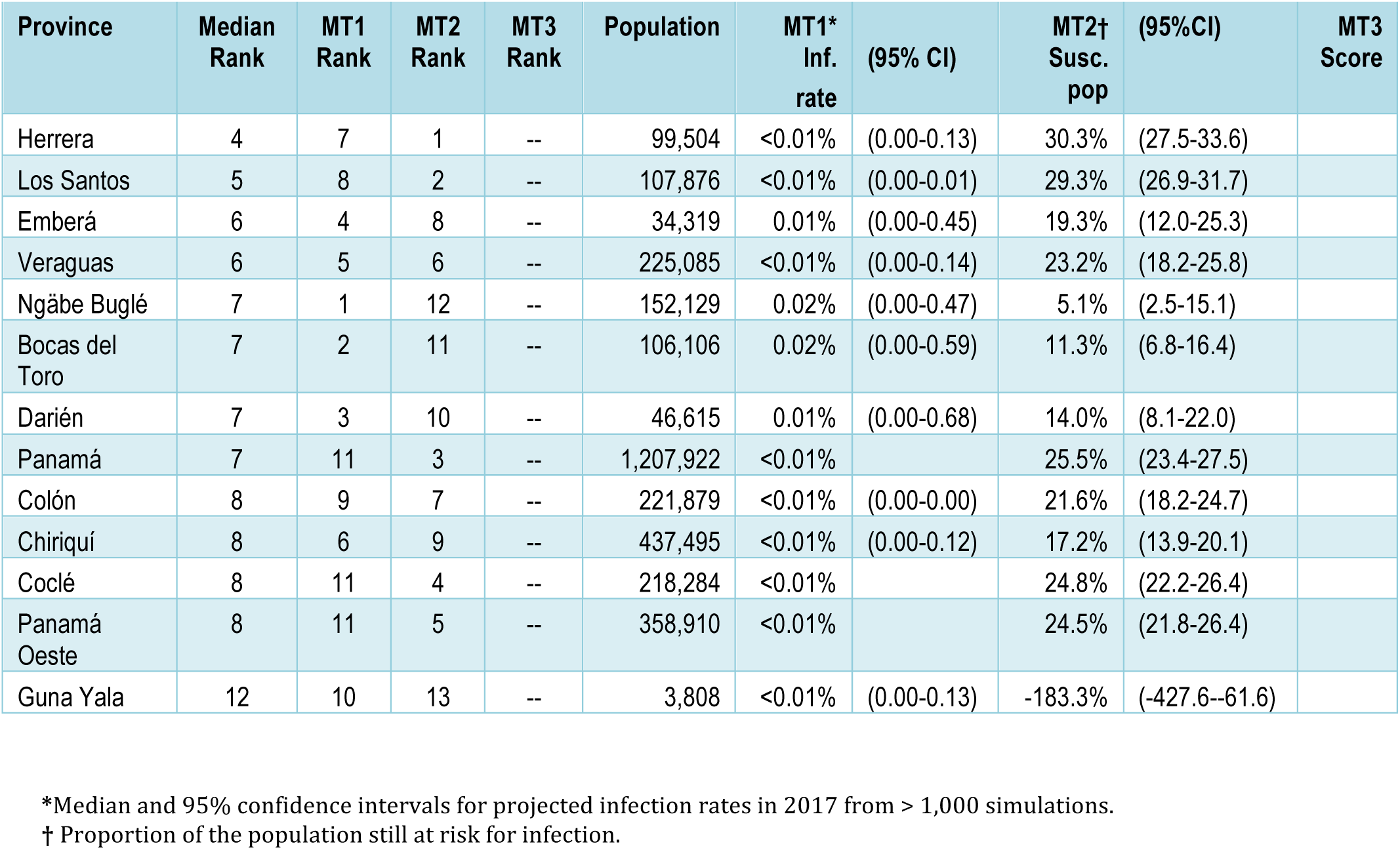
Panama results by province

**Table 8:**
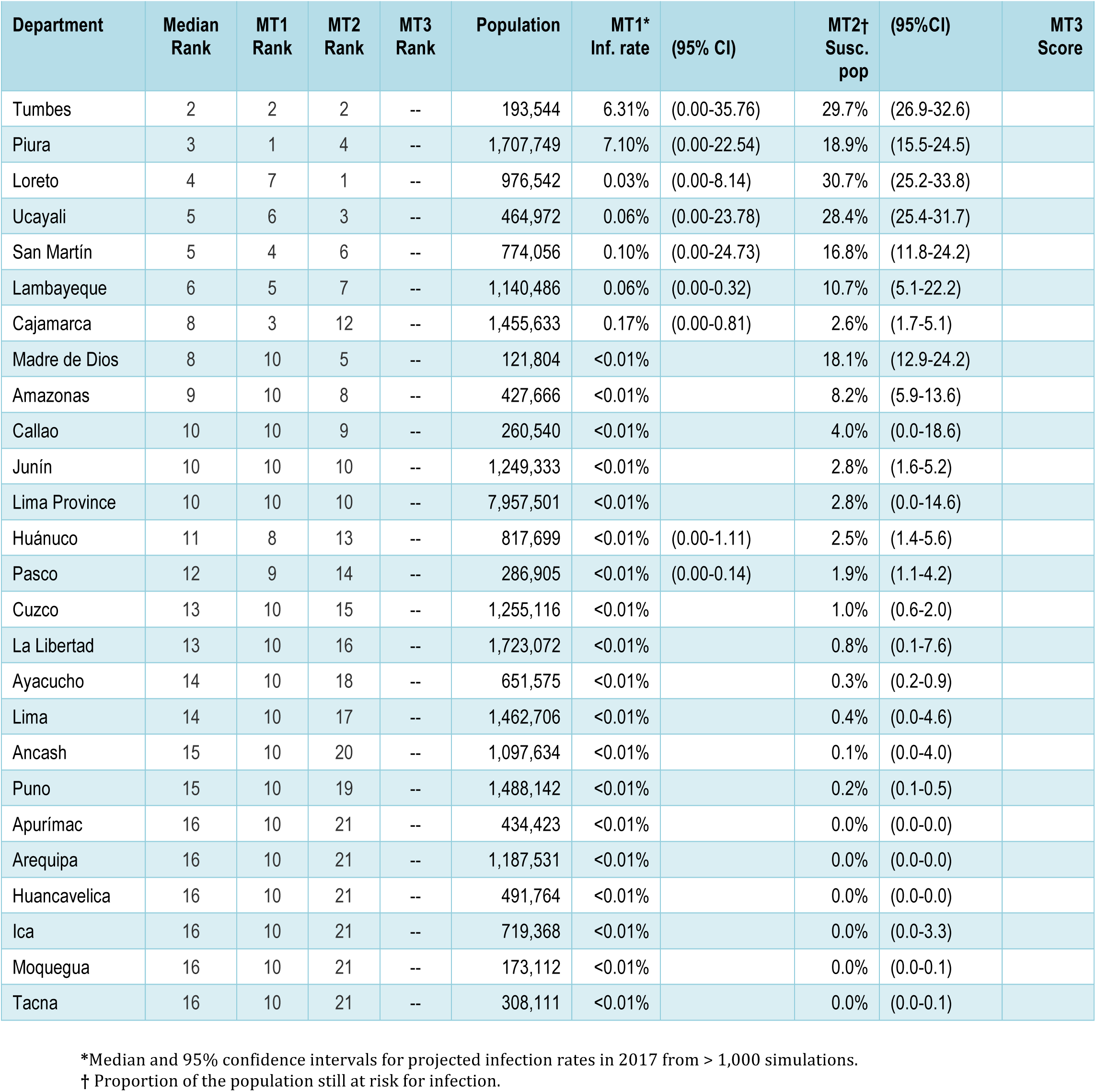
Peru modeling results by department

| Department | Median Rank | MT1 Rank | MT2 Rank | MT3 Rank | Population | MT1* Inf. rate | (95% CI) | MT2† Susc. pop | (95%CI) | MT3 Score |
| --- | --- | --- | --- | --- | --- | --- | --- | --- | --- | --- |
| Tumbes | 2 | 2 | 2 | -- | 193,544 | 6.31% | (0.00-35.76) | 29.7% | (26.9-32.6) |  |
| Piura | 3 | 1 | 4 | -- | 1,707,749 | 7.10% | (0.00-22.54) | 18.9% | (15.5-24.5) |  |
| Loreto | 4 | 7 | 1 | -- | 976,542 | 0.03% | (0.00-8.14) | 30.7% | (25.2-33.8) |  |
| Ucayali | 5 | 6 | 3 | -- | 464,972 | 0.06% | (0.00-23.78) | 28.4% | (25.4-31.7) |  |
| San Martín | 5 | 4 | 6 | -- | 774,056 | 0.10% | (0.00-24.73) | 16.8% | (11.8-24.2) |  |
| Lambayeque | 6 | 5 | 7 | -- | 1,140,486 | 0.06% | (0.00-0.32) | 10.7% | (5.1-22.2) |  |
| Cajamarca | 8 | 3 | 12 | -- | 1,455,633 | 0.17% | (0.00-0.81) | 2.6% | (1.7-5.1) |  |
| Madre de Dios | 8 | 10 | 5 | -- | 121,804 | <0.01% |  | 18.1% | (12.9-24.2) |  |
| Amazonas | 9 | 10 | 8 | -- | 427,666 | <0.01% |  | 8.2% | (5.9-13.6) |  |
| Callao | 10 | 10 | 9 | -- | 260,540 | <0.01% |  | 4.0% | (0.0-18.6) |  |
| Junín | 10 | 10 | 10 | -- | 1,249,333 | <0.01% |  | 2.8% | (1.6-5.2) |  |
| Lima Province | 10 | 10 | 10 | -- | 7,957,501 | <0.01% |  | 2.8% | (0.0-14.6) |  |
| Huánuco | 11 | 8 | 13 | -- | 817,699 | <0.01% | (0.00-1.11) | 2.5% | (1.4-5.6) |  |
| Pasco | 12 | 9 | 14 | -- | 286,905 | <0.01% | (0.00-0.14) | 1.9% | (1.1-4.2) |  |
| Cuzco | 13 | 10 | 15 | -- | 1,255,116 | <0.01% |  | 1.0% | (0.6-2.0) |  |
| La Libertad | 13 | 10 | 16 | -- | 1,723,072 | <0.01% |  | 0.8% | (0.1-7.6) |  |
| Ayacucho | 14 | 10 | 18 | -- | 651,575 | <0.01% |  | 0.3% | (0.2-0.9) |  |
| Lima | 14 | 10 | 17 | -- | 1,462,706 | <0.01% |  | 0.4% | (0.0-4.6) |  |
| Ancash | 15 | 10 | 20 | -- | 1,097,634 | <0.01% |  | 0.1% | (0.0-4.0) |  |
| Puno | 15 | 10 | 19 | -- | 1,488,142 | <0.01% |  | 0.2% | (0.1-0.5) |  |
| Apurímac | 16 | 10 | 21 | -- | 434,423 | <0.01% |  | 0.0% | (0.0-0.0) |  |
| Arequipa | 16 | 10 | 21 | -- | 1,187,531 | <0.01% |  | 0.0% | (0.0-0.0) |  |
| Huancavelica | 16 | 10 | 21 | -- | 491,764 | <0.01% |  | 0.0% | (0.0-0.0) |  |
| Ica | 16 | 10 | 21 | -- | 719,368 | <0.01% |  | 0.0% | (0.0-3.3) |  |
| Moquegua | 16 | 10 | 21 | -- | 173,112 | <0.01% |  | 0.0% | (0.0-0.1) |  |
| Tacna | 16 | 10 | 21 | -- | 308,111 | <0.01% |  | 0.0% | (0.0-0.1) |  |

\*Median and 95% confidence intervals for projected infection rates in 2017 from > 1,000 simulations.
† Proportion of the population still at risk for infection.

**Table 9:** Probability of projected annual Zika virus infection rates ≥ 10% in 2017 for municipalities in the Americas

| Country | State | Municipality | Municipality population | Model 1<br>Probability of<br>infection rate $\geq 10\%$<br>in 2017 | (95% CI) |
| --- | --- | --- | --- | --- | --- |
| Ecuador | Sucumbios | Lago Agrio/Nueva Loja | 249,959 | 49% | (46-52) |
| Peru | Tumbes | Tumbes | 283,424 | 48% | (45-51) |
| Peru | Piura | Piura | 1,730,114 | 45% | (43-48) |
| Mexico | Sinaloa | Los Mochis | 1,049,654 | 40% | (38-43) |
| Bolivia | Gran Chaco | Yacuiba | 243,077 | 27% | (24-29) |
| Ecuador | Orellana | El Coca | 151,629 | 21% | (19-24) |
| Paraguay | Alto Parana | Ciudad del Este | 2,336,625 | 21% | (19-23) |
| Mexico | Tamaulipas | Tampico | 1,835,390 | 18% | (16-20) |
| Honduras | Gracias a Dios | Puerto Lempira | 106,834 | 16% | (14-18) |
| Ecuador | Esmeraldas | Esmeraldas | 599,629 | 12% | (10-14) |
| Peru | San Martin | Tarapoto | 1,397,529 | 10% | (9-12) |
| Colombia | Narino | Tumaco | 275,111 | 9% | (7-10) |
| Peru | Ucayali | Pucallpa | 694,810 | 8% | (7-10) |
| Mexico | Colima | Colima | 957,465 | 8% | (6-9) |
| Mexico | Baja California Sur | Cabo San Jose | 149,391 | 8% | (6-9) |
| Honduras | Colon | Guanaja/Tocoa | 221,233 | 6% | (5-8) |
| Colombia | Guainia | Puerto Inirida | 53,718 | 6% | (5-8) |
| Mexico | Michoacan | Lazaro Cardenas | 341,119 | 6% | (4-7) |
| Mexico | Sinaloa | Culiacan | 1,481,563 | 5% | (4-7) |
| Colombia | Vichada | Puerto Carreno | 39,127 | 5% | (4-6) |
| Venezuela | Bolivar | Santa Elena | 8,160 | 5% | (3-6) |
| Mexico | Oaxaca | Santa Maria Huatulco | 665,936 | 4% | (3-6) |
| Mexico | Oaxaca | Puerto Escondido | 464,002 | 4% | (3-5) |
| Mexico | Colima | Manzanillo | 390,234 | 4% | (3-5) |
| Mexico | Chiapas | Tapachula | 921,029 | 3% | (2-4) |
| Mexico | Nayarit | Tepic | 934,532 | 3% | (2-4) |
| Mexico | Sinaloa | Mazatlan | 699,002 | 3% | (2-4) |
| Brazil | Parana | Umuarama | 518,546 | 3% | (2-4) |
| Mexico | Veracruz | Minatitlan | 1,881,784 | 3% | (2-4) |
| Colombia | Amazonas | Tarapaca | 50,271 | 2% | (2-3) |

This modeling approach has been used previously to estimate the transmission and spread of pandemic influenza and Ebola [Tizzoni 2012; Gomes 2014; Poletto 2014]. In order to validate the approach for Zika virus, the authors compared model-based projections to independent surveillance reports of numbers of infections in Colombia, microcephaly cases in Brazil, and travel-associated disease cases in the continental United States and Europe [Zhang 2017].

### Modeling Team 2 (MT2)^2^

This approach compares model projections of infection rates with estimates of cumulative infections to date. Locations with a high projected infection rate that have experienced low transmission to date are presumed to be good candidates for vaccine trials because a relatively large portion of the population is still susceptible and likely to become infected before the epidemic subsides. The infection rate projections are informed by spatial layers of variables pertaining to human demography, purchasing power parity, temperature, and vector occurrence probability. Relationships between these variables and infection rates are drawn from the theoretical and empirical literature on other pathogens transmitted by *Aedes aegypti* mosquitoes, and remaining uncertainties in the model’s form are calibrated to seroprevalence estimates following introduction of chikungunya or Zika virus in immunologically naïve populations. The result is a spatial layer of location-specific projections of the number of Zika virus infections that are expected to occur in each 5 km by 5 km area across Latin America and the Caribbean [Perkins 2016]. The timeframe for the projected infections is from the beginning of the epidemic until however long it takes for the epidemic to end due to the buildup of sufficient herd immunity. The extent of herd immunity that is sufficient to end the epidemic is positively associated with transmission potential (i.e., a greater proportion of the population must build-up immunity in settings with intense transmission). Although this model does not predict the precise timeframe over which the Zika virus epidemic will run its course, other estimates suggest local epidemics may be extinguished by herd immunity 2–3 years after the initial introduction of Zika virus [Ferguson 2016].

Estimates of cumulative infections to date are based on a combination of cumulative reported cases and assumptions about the proportion of infections that are reported, denoted as *p*. It is generally accepted that *p* is extremely variable across settings and difficult to ascertain. Based on discussions among the modeling teams, there is general agreement that *p* may often be around 1-2%, could sometimes be as high as 5%, and is unlikely to exceed 10% in the settings under consideration. Given this overall uncertainty about *p* and collective opinion about what values it may likely take, we assumed that *p* ∼ 0.01 + 0.9Beta(1.2,5). This implies that the expected value of *p* is 0.027 and that it does not exceed 0.1 or fall below 0.01. This approach to parameterizing *p* is similar to formulating a prior probability distribution based on “expert opinion” in a Bayesian analysis. For context, we note a limited number of published estimates of *p*: 0.015 (95% CI: 0.036-0.022) on Yap Island in 2007 [Duffy 2009]; 0.115 (95% CI: 0.073-0.179) in French Polynesia in 2013-2014 [Kucharski 2016]; 0.021 (95% CI: 0.017-0.025) in Puerto Rico in 2016 [Chevalier 2017]; and 0.010 (standard deviation = 0.0093) across the Americas as a whole in 2013-2016 [Zhang 2017]. Subsequent refinements of this approach will seek to incorporate additional estimates and to more formally characterize uncertainty about *p*.

The authors then compare the projected number of infections that will occur before the first wave of the epidemic concludes to estimates of the current cumulative incidence of infection to estimate the proportion of the population that remains at risk for infection. Locations with a large discrepancy between numbers of projected total infections and estimated infections to date are interpreted to be good candidates for vaccine trial sites (**Tables 1–8**).

### Modeling Team 3 (MT3)^3^

This approach uses the age-specific incidence of dengue to calculate the associated force of infection for dengue per administrative unit [Cummings 2009; Ferguson 2016; Rodriguez-Barraquer 2016]. This hazard of infection previously was shown to correlate with Zika incidence in Colombia and with microcephaly incidence in Brazil. In each country, the relationship between dengue force of infection and reported Zika virus disease incidence (Mexico, Colombia) or microcephaly incidence (Brazil) is calculated based on a presumed linear relationship between the square root of Zika incidence (or a proxy) and the force of infection. A statistical probability score per administrative unit is calculated and is the probability of seeing the number of observed cases or greater for a given force of infection, if the square root of incidence is normally distributed with the predicted mean and observed variance of the residuals. This score is used to rank areas, and can be roughly interpreted as the probability of seeing the observed number of cases or fewer if the Zika epidemic has completed in an area with that force of infection for dengue (**Tables 1–8**).

### Integrating the model results

Each province, state, or department in the eight priority countries was ranked according to the primary outcome measure for each of the models that provided data for that country (i.e., all three models for Brazil, Colombia, and Mexico, and models 1 and 2 for Costa Rica, Dominican Republic, Ecuador, Panama, and Peru). The MT3 output was not provided for countries for which age-specific dengue incidence data was unavailable.

The median rank for the available models was calculated and states/provinces were ordered and mapped for each country. The consistency among models was assessed by identifying in each country the states, provinces, or departments ranking within the top quartile by two or more models.

We used data from MT1 to identify states, provinces, or departments with a median projected infection rate ≥10% to approximate a symptomatic infection rate ≥2% in the eight priority countries, assuming that roughly 20% of cases are symptomatic [Duffy 2009]. We also used Model 1 data to identify municipalities in any country/territory in the Americas with ≥5% probability of having a projected Zika virus infection rate ≥10% in 2017.

## Results

In order to compare the modeling results, we provide a list of locations (state, province or department) for the eight priority countries, prioritized by the median rank of the models’ outcomes. In **Tables 1–8** and **Figures 1–8** we provide the modeling results of the different models. We show the ranking of the different locations across models and compute a median rank. In addition, we provide the original results of the different models from which we construct the ranking of locations.

**Figure 1:**
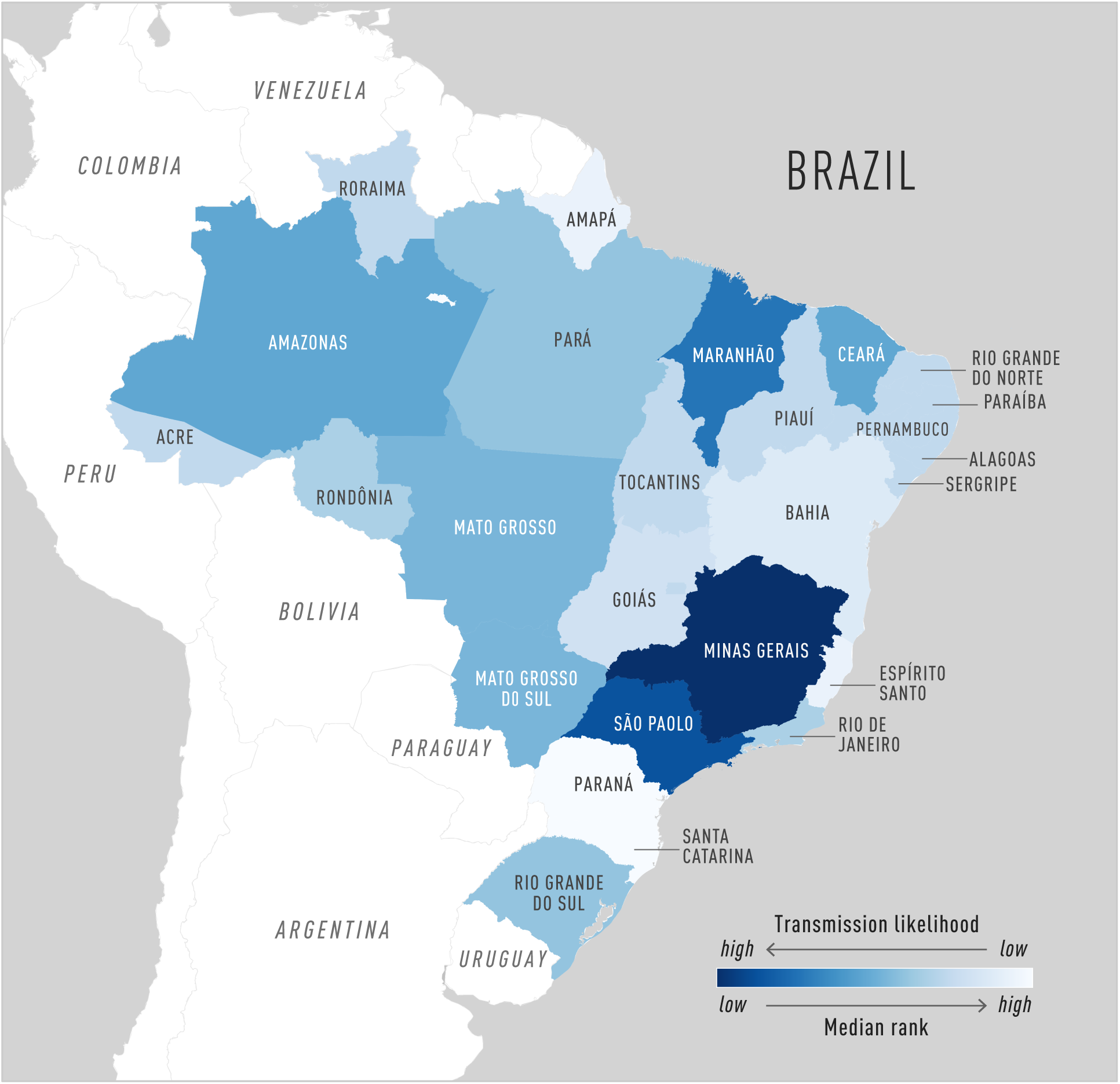
Median rank for the three models for each state in Brazil. The color scheme shows how each state ranks within the country. That is, the lower the rank (dark blue), the higher the relative likelihood of future Zika transmission.

**Figure 2:**
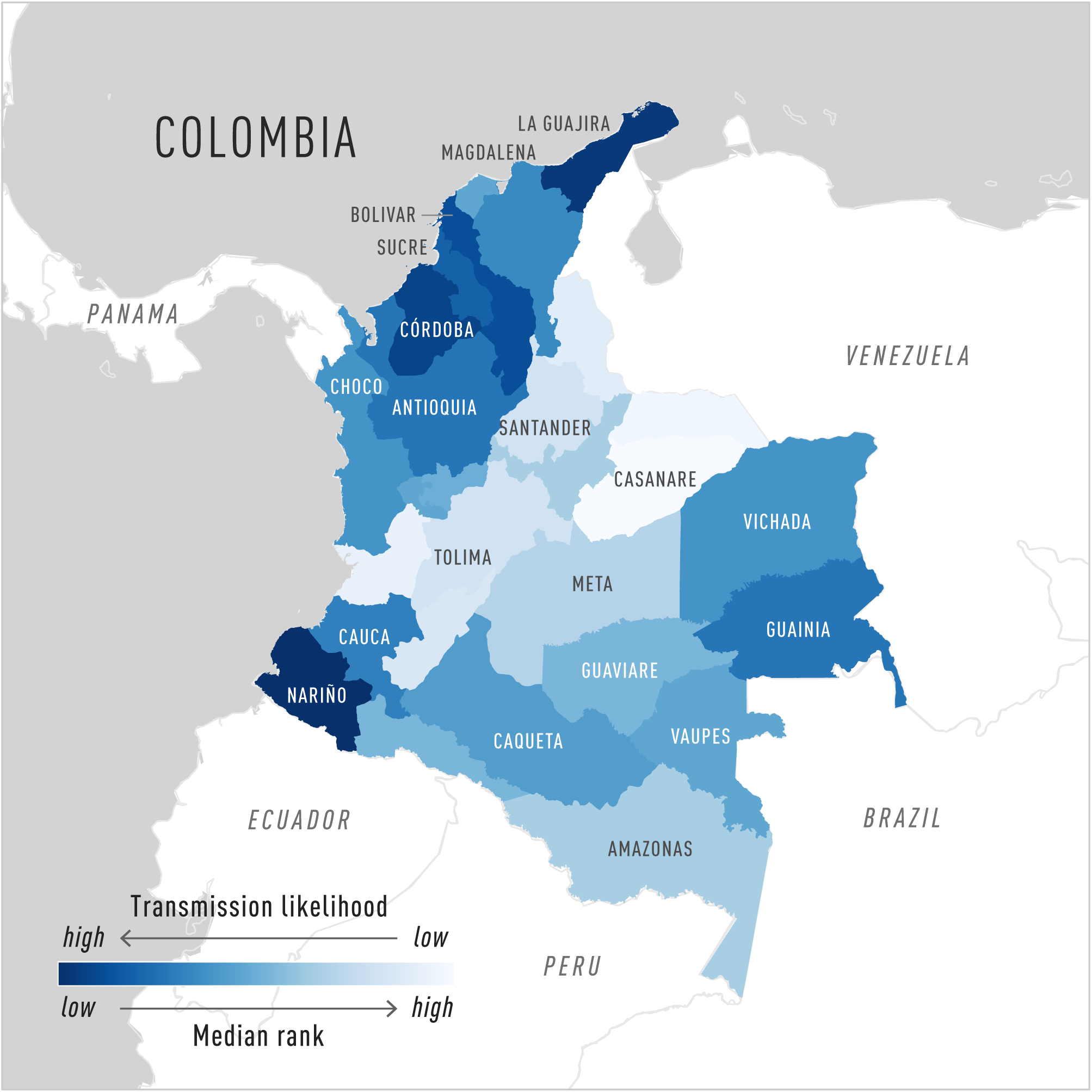
Median rank for the three models for each state in Colombia. The color scheme shows how each state ranks within the country. That is, the lower the rank (dark blue), the higher the relative likelihood of future Zika transmission.

**Figure 3:**
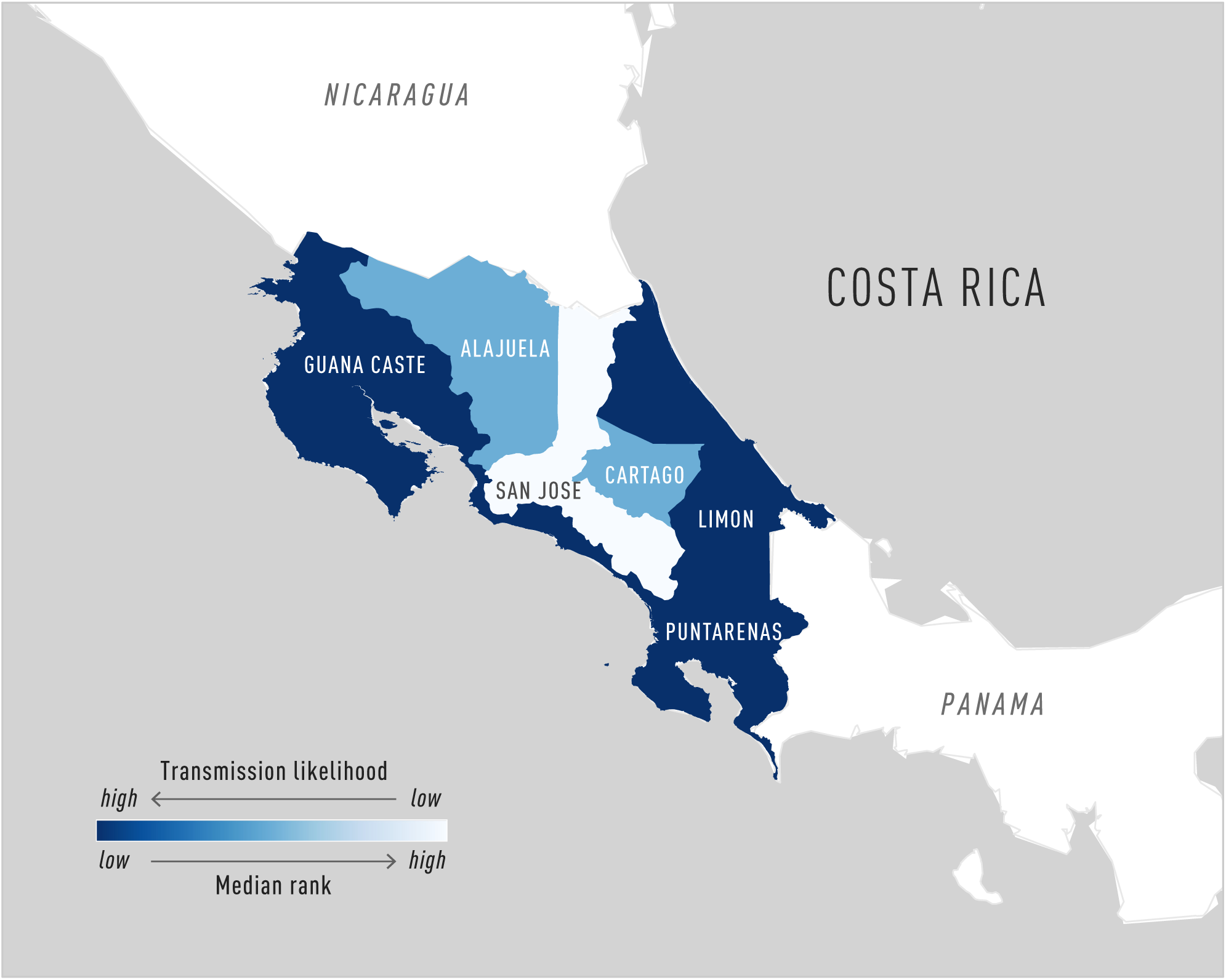
Median rank for the three models for each province in Costa Rica. The color scheme shows how each state ranks within the country. That is, the lower the rank (dark blue), the higher the relative likelihood of future Zika transmission.

**Figure 4:**
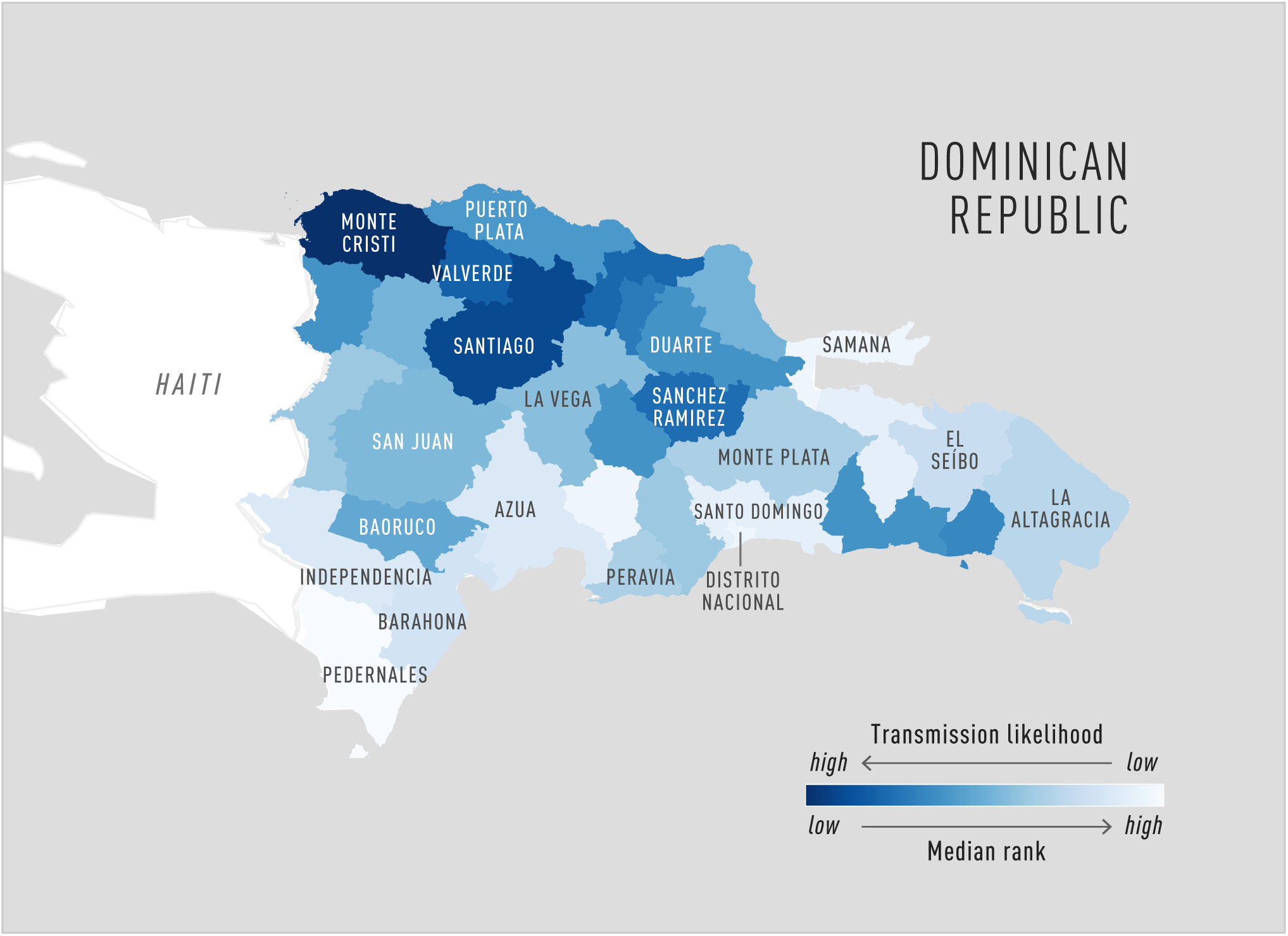
Median rank for the three models for each province in Dominican Republic. The color scheme shows how each state ranks within the country. That is, the lower the rank (dark blue), the higher the relative likelihood of future Zika transmission.

**Figure 5:**
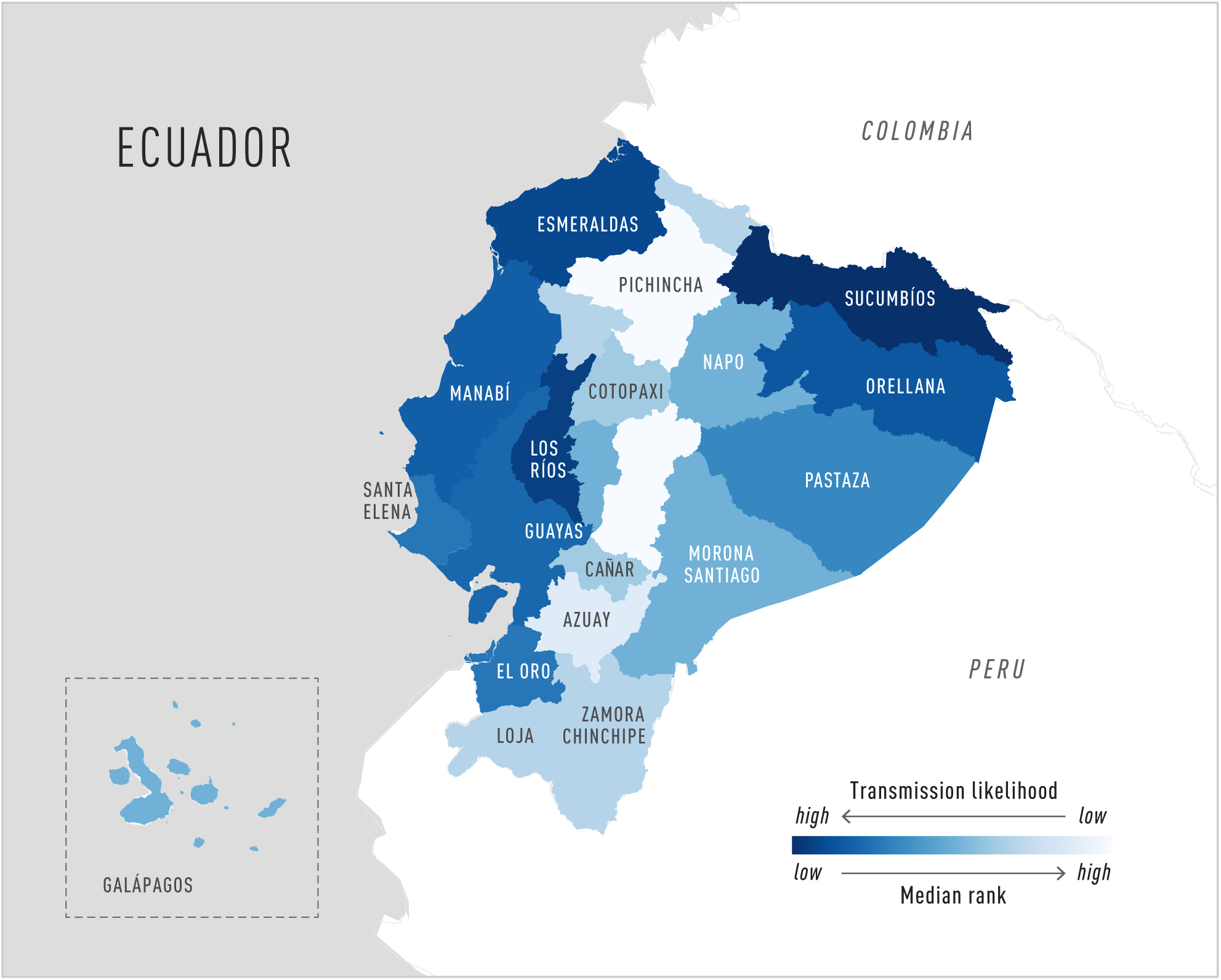
Median rank for the three models for each province in Ecuador. The color scheme shows how each state ranks within the country. That is, the lower the rank (dark blue), the higher the relative likelihood of future Zika transmission.

**Figure 6:**
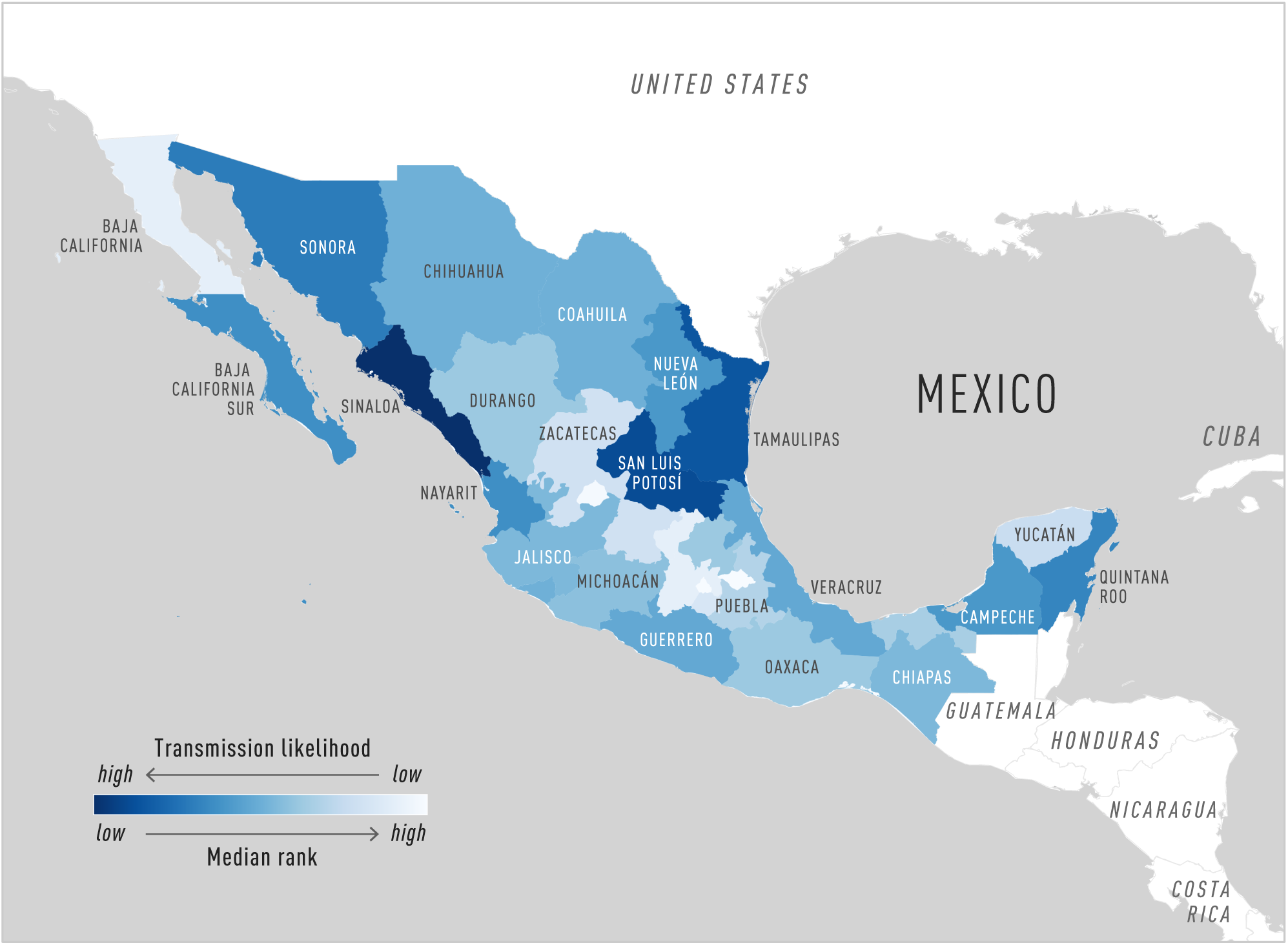
Median rank for the three models for each state in Mexico. The color scheme shows how each state ranks within the country. That is, the lower the rank (dark blue), the higher the relative likelihood of future Zika transmission.

**Figure 7:**
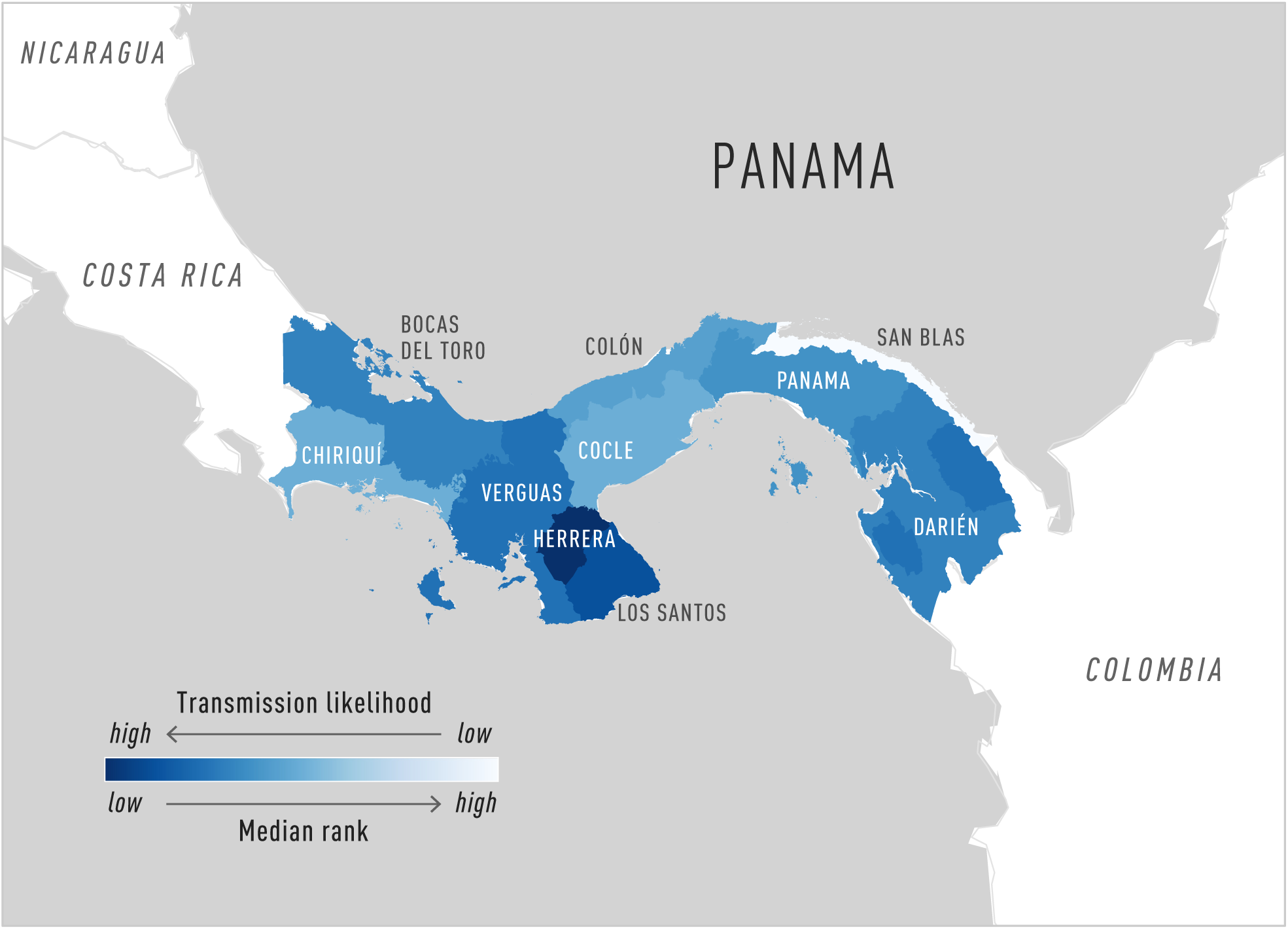
Median rank for the three models for each province in Panama. The color scheme shows how each state ranks within the country. That is, the lower the rank (dark blue), the higher the relative likelihood of future Zika transmission.

**Figure 8:**
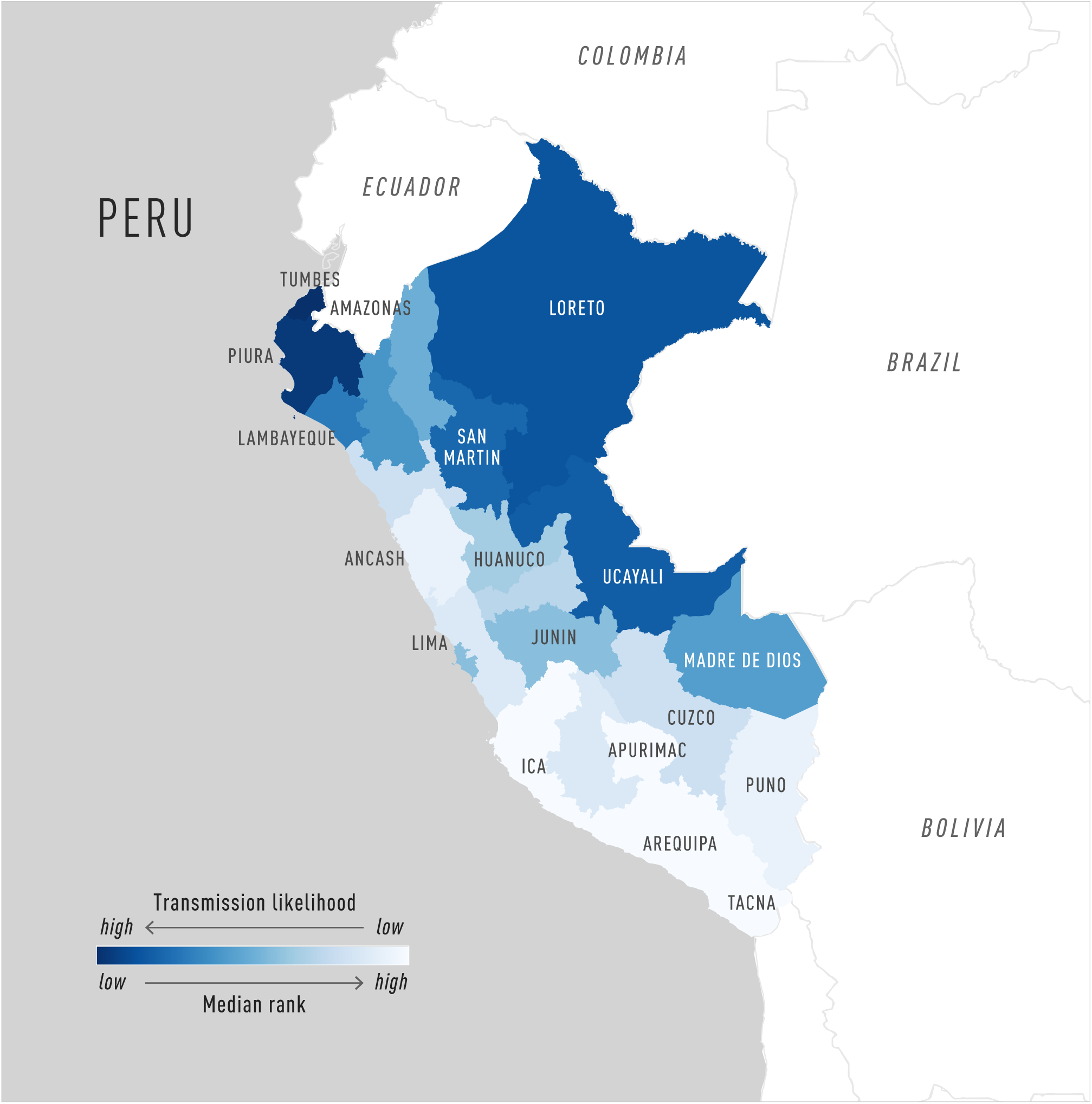
Median rank for the three models for each department in Peru. The color scheme shows how each state ranks within the country. That is, the lower the rank (dark blue), the higher the relative likelihood of future Zika transmission.

The results show that 18 locations (states, provinces, or departments) in six different countries ranked within the top quartile for the given country by two or more models. These locations are the following: Minas Gerais, São Paulo, and Maranhão states in Brazil; Nariño, Guajira, Córdoba, Bolívar, and Sucre states in Colombia; Monte Cristi, and Santiago provinces in Dominican Republic; Sucumbios and Los Ríos provinces in Ecuador; Sinaloa, San Luis Potosí, and Tamaulipas states in Mexico; Tumbes, Piura, Ucayali, and San Martin departments in Peru. For Costa Rica and Panama there are no locations ranked within the top quartile by two models. The median rank and range of model ranks (black line) for each state/province within each country are shown in **Figure 9**. The three models generate a relative ranking in each country; the ranking does not necessarily reflect an absolute risk or infer high virus transmission activity in the top-ranked locations. Locations may have low expected Zika transmission activity but still be ranked in the top places when compared to other places within the same country that show even lower activity.

**Figure 9:**
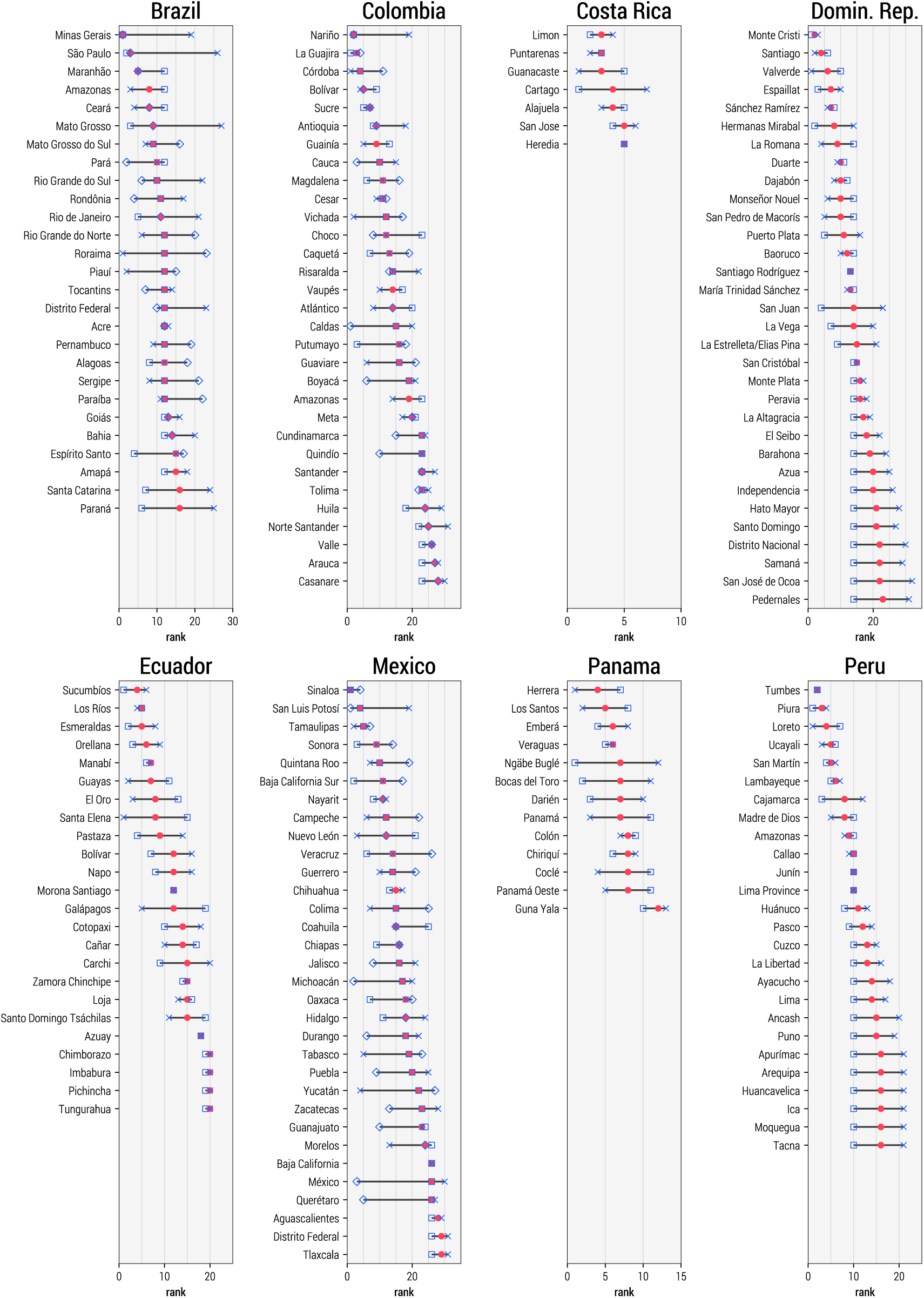
Median rank (red dot) and range of model ranks (black line) for each state/province within each country. Individual models are indicated by open blue shapes (square = Model 1; cross = Model 2; diamond = Model 3). Additional details can be found in Tables 1–8.

Only five locations in two countries have a projected median Zika virus infection rate larger than 5% in 2017: Sucumbios, Esmeraldas, and Orellana provinces in Ecuador; and Tumbes and Piura departments in Peru (see **Tables 1–8**). When comparing across the three models, three of the previous five locations with a projected Zika virus infection rate larger than 5%, ranked within the top quartile for their country by two or more of the models: Sucumbios in Ecuador and Tumbes and Piura in Peru.

MT1 also offers the possibility of zooming in the different locations, allowing the study of the outbreak at the level of municipalities or urban areas. Through this analysis, we observe that 21 municipalities have a probability larger than 5% of having a projected Zika virus infection rate of at least 10% in 2017 (see **Table 9**). From the municipalities identified by MT1, nine of them are located within regions that are also identified by two or more of the models. The municipalities are the following:

- Colombia: Tumaco in Nariño state.
- Ecuador: Lago Agrio/Nueva Loja in Sucumbios province.
- Mexico: Los Mochis and Culiacan in Sinaloa state, and Tampico in Tamaulipas state.
- Peru: Piura in Piura department, Tumbes in Tumbes department, Tarapoto in San Martín department, and Pacallpa in Ucayali department.

In **Figures 1–8,** we provide a geographical visualization of the administrative units ranked in the tables. We use a color map associated to the rank order to localize places according to their likelihood of Zika transmission. The purpose of the maps is to illustrate any potential regional clustering of provinces/states with relatively higher likelihood of activity within each country. We again stress that the maps report the median ranking, as obtained by aggregating the results of the three models, and that the ranking is just indicative of the relative likelihood of future transmission within each country.

## Summary

These preliminary findings provide states/provinces and municipalities in eight priority countries where study sites may have increased likelihood of having sufficient Zika virus transmission to meet the efficacy end points in 2017. Due to substantial differences and uncertainties in data between countries, we limited the comparisons to estimates for subnational areas within each of the priority countries.

All of the evaluated subnational areas in the priority countries had low projected incidence rates in 2017. Only three provinces or departments in two countries had a projected Zika virus infection rate >5% and ranked within the top quartile for their country by two or more of the models. We also identified relatively few municipalities that have a projected Zika virus infection rate ≥10% and are located in states with consistently high rankings by two or more of the models.

In summary, the models suggest that the total number of participants, number of study sites, and/or duration of study follow-up may need to be increased to meet the efficacy end points. The findings also support initiating a high number of study sites in multiple geographic areas to maximize the likelihood of having study capacity in one or more areas that experience Zika virus infections in 2017 and provide flexibility to responsively increase enrollment in areas with the highest incidence of infection

This report is made available to share the approach and preliminary findings with the research community. Results should be interpreted cautiously given the model limitations and assumptions. Furthermore, projecting the Zika virus transmission at seasonal and longer timescales increases uncertainty, especially given the lack of comprehensive, quality surveillance data on current and previous Zika virus transmission activity. The modeling teams are continuing these efforts and will provide an updated report which will incorporate: 1) refined modeling methods, 2) updated surveillance data, and 3) further integration and discussion of similarities and differences between the model findings.

## Footnotes

1 MT1 includes Alessandro Vespignani, Ana Pastore y Piontti, Kaiyuan Sun, Matteo Chinazzi, M. Elizabeth Halloran, Ira M. Longini, Stefano Merler, Luca Rossi, and Qian Zhang.

2 MT2 includes Alex Perkins, Amir Siraj, Christopher Barker, and Robert Reiner.

3 MT3 includes Justin Lessler, Isabel Rodriguez-Barraquer, Derek Cummings, and Neil Ferguson.

